# Minimal Positional Substring Cover: A Haplotype Threading Alternative to Li & Stephens Model

**DOI:** 10.1101/2023.01.04.522803

**Authors:** Ahsan Sanaullah, Degui Zhi, Shaojie Zhang

## Abstract

The Li & Stephens (LS) hidden Markov model (HMM) models the process of reconstructing a haplotype as a mosaic copy of haplotypes in a reference panel (haplotype threading). For small panels the probabilistic parameterization of LS enables modeling the uncertainties of such mosaics, and has been the foundational model for haplotype phasing and imputation. However, LS becomes inefficient when sample size is large (tens of thousands to millions), because of its linear time complexity (*O*(*MN*), where *M* is the number of haplotypes and *N* is the number of sites in the panel). Recently the PBWT, an efficient data structure capturing the local haplotype matching among haplotypes, was proposed to offer fast methods for giving some optimal solution (Viterbi) to the LS HMM. But the solution space of the LS for large panels is still elusive. Previously we introduced the Minimal Positional Substring Cover (MPSC) problem as an alternative formulation of LS whose objective is to cover a query haplotype by a minimum number of segments from haplotypes in a reference panel. The MPSC formulation allows the generation of a haplotype threading in time constant to sample size (*O*(*N*)). This allows haplotype threading on very large biobank scale panels on which the LS model is infeasible. Here we present new results on the solution space of the MPSC by first identifying a property that any MPSC will have a set of required regions, and then proposing a MPSC graph. In addition, we derived a number of optimal algorithms for MPSC, including solution enumerations, the Length Maximal MPSC, and *h*-MPSC solutions. In doing so, our algorithms reveal the solution space of LS for large panels. Even though we only solved an extreme case of LS where the emission probability is 0, our algorithms can be made more robust by PBWT smoothing. We show that our method is informative in terms of revealing the characteristics of biobank-scale data sets and can improve genotype imputation.

## 1 Introduction

Human chromosomes are a mosaic of ancestral chromosomal segments, resulting from accumulated recombination events. By way of transitivity, when querying a chromosomal haplotype against a panel of haplotypes, one can view the query haplotype as a mosaic of identical-by-descent (IBD) segments with haplotypes in the panel. Such mosaics, also called haplotype threadings, are the foundational model for haplotype phasing, genotype imputation, and relatedness inference.

For many years, the state of the art haplotype threading method was the Li & Stephens (LS) Model [7]. The Li & Stephens model produces a haplotype threading by using a Hidden Markov Model (HMM) where penalties are incurred for mismatches and switching between different haplotypes in adjacent sites the panel in the threading. When sample size is small, the matching segments, i.e., the continuous streak of match with the same haplotype template in the panel, are typically short, and the Li & Stephens model can be parameterized to achieve a balance between mismatch tolerance and minimizing switching. Indeed, most current phasing and imputation methods are based on the Li & Stephens model [1, 2, 4, 5, 8, 12].

However, when sample size of the panel is large, Li & Stephens model becomes inefficient, as its time complexity is linear to the size of the panel, *O*(*MN*), where *M* is the number of haplotypes in the panel and N is the number of sites per haplotype. The latest imputation methods based on LS have used PBWT to quickly identify a subset of templates as “surrogate parents” of the query, but still use LS HMM to sample the threading over the subset [2, 12].

Recent work has attempted to obtain the optimal haplotype threading in the Li & Stephens model in time sublinear to the size of the panel. Gerton Lunter described “fastLS”, an algorithm that implements the Li & Stephens model and obtains the optimal haplotype threading through the use of the Burrows-Wheeler transform [9]. His algorithm obtains the optimal haplotype threading orders of magnitude faster than the Viterbi algorithm. Yohei Rosen and Benedict Paten also described an algorithm that achieved runtime orders of magnitude faster than the Viterbi algorithm [11]. Their method achieves this using the efficient sparse representation of haplotypes and the lazy evaluation of dynamic programming. These algorithms have also been claimed to have runtime sublinear to the size of the reference panel. However, this claim has only been shown empirically.

In this work, we discuss an alternative formulation of the haplotype threading problem. This is based on the observation that when the size of the panel is large enough, approaching the size of the population, the IBD segments shared between the query and the template haplotypes can be much longer. Also, the error rates on modern large panels are very low (~0.1%). Therefore, the problem of haplotype threading for large panels is Li & Stephens model at the regime where the mismatch rates and switch rate are very low. In other words, it is of interest to make combinatorial formulations of the problem.

In a previous work, Sanaullah et al. introduced a new formulation of haplotype threading, the Minimal Positional Substring Cover (MPSC) [13]. The MPSC problem is, given a query haplotype *z* and a set of haplotypes *X*, find a smallest set of segments of haplotypes in *X* that covers *z*. This formulation corresponds to LS model with 0 mismatch rate and the only objective is to minimize the switch rate. Obviously, the MPSC formulation is too limited as it does not tolerate mismatches. In a sense, MPSC is a more general formulation of haplotype threading. The traditional haplotype threading is a special case of MPSC with non-overlapping segments. MPSC formulation captures the fact that while the switching of templates indicates some recombination events, the exact breakpoint of the recombination events may be anywhere within the overlapping region between the templates. Furthermore, this combinatorial formulation is theoretically attractive because it is can be solved in worst case *O*(*N*) time [13]. I.E. given a PBWT of the reference panel, a MPSC haplotype threading of a query haplotype can be done in time independent to the number of haplotypes in the reference panel. Further, a number of variations of the MPSC formulation can be solved efficiently, including Leftmost and Rightmost MPSCs, MPSC composed of only set maximal matches, and *h*-MPSC.

In this work, we expand upon the abilities of the MPSC formulation of haplotype threading greatly. Our major contributions are the exploration of the solution space of minimal positional substring covers, the description and solving of a new variation of the MPSC problem, the Length Maximal MPSC, the providing of an improved algorithm for the *h*-MPSC problem, and the demonstration of usefulness of the MPSC formulation haplotype threading through an imputation benchmark. Table 1 summarizes the major algorithmic contributions of this paper.

**Table 1.**
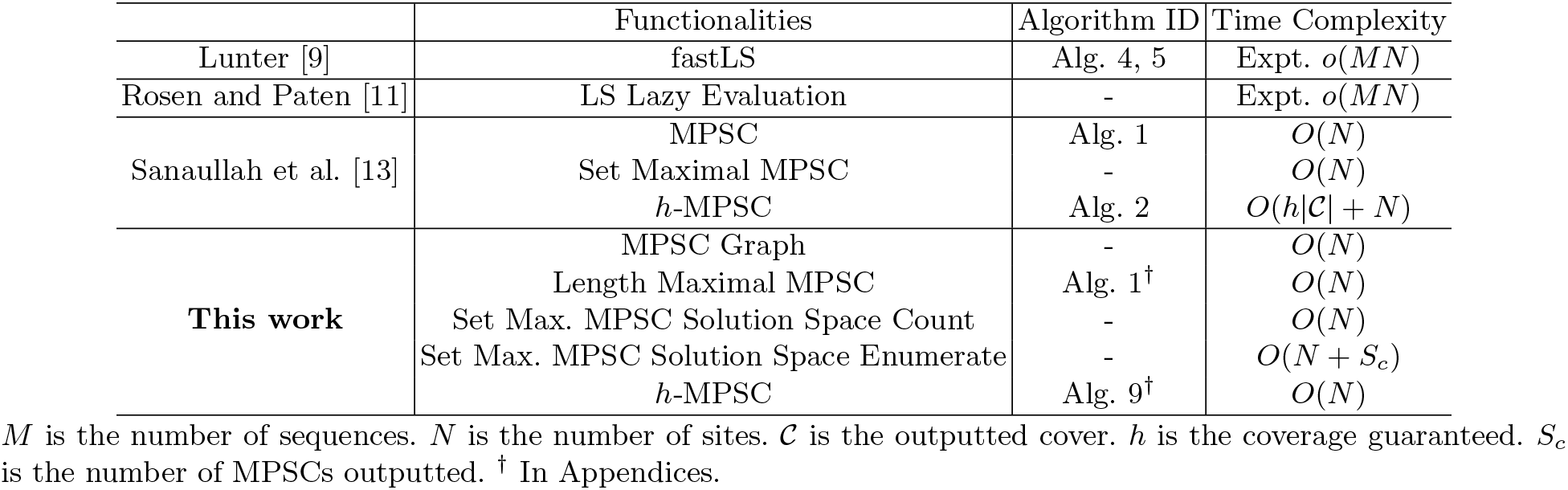
Summary of algorithms on haplotype threading.

## 2 Background

By following the definitions in [13], a string *z* with *N* characters is indexed from 0 to *N* – 1. The first character of *z* is *z*[0] and the last is *z*[*N* – 1]. The string *z*[*i, j*] is the substring of *z* that starts at character *i* and ends at character *j*.

A **positional substring** of a string *z* is a 3-tuple, (*i, j, z*), where *i* and *j* are nonegative integers, *i* ≤ *j* + 1 ≤ |*z*|, and *z* is the “source” of the substring. The substring corresponding to the positional substring (*i, j, z*) is *z*[*i, j*]. If *i* = *j* + 1, then (*i, j, z*) corresponds to the empty string, *ϵ*. A non-empty positional substring (*i, j, z*) is contained in a string *s* if 0 ≤ *i* ≤ *j* < |*s*| and *s*[*i, j*] = *z*[*i, j*]. An empty positional substring (*i, i* −1, *z*) is contained in a string *s* iff 0 ≤ *i* ≤ |*s*|. Two positional substrings, (*i, j, s*) and (*k, l, t*) are equal iff *i* = *k, j* = l, and *s*[*i, j*] = *t*[*k, l*]. The length of a positional substring (*i, j, s*) is *j* −*i* + 1.

A **positional substring cover**, 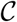, of a string *z* by a set of strings *X* is a set of positional substrings such that every character of *z* is contained in a positional substring and every positional substring in the set is present in *z* and a string in *X*. The “source,” *s*, of a positional substring (*i, j, s*) in a positional substring cover can be any string *s* s.t. *s*[*i, j*] = *x*[*i, j*] = *z*[*i, j*] for some *x* ∈ *X*. The size of a positional substring cover is the number of positional substrings it contains. The length of a positional substring cover is the sum of the lengths of its positional substrings.

*π* is a projection on tuples. *π*_1_(*i, j, z*) = *i*, *π*_2_(*i, j, z*) = *j*, and *π*_3_(*i, j, z*) = *z*.

### 2.1 PBWT

The algorithms described in this paper require a prebuilt positional Burrows-Wheeler transform. The positional Burrows-Wheeler transform (PBWT) is a data structure that allows memory efficient representation of a set of binary haplotypes of the same length [6]. The PBWT also allows time efficient outputting of matches between haplotypes in the panel. Here we provide a brief description of the PBWT and its algorithms.

The PBWT stores *X*, a set of *M* binary strings of length *N*. It also stores four arrays of size roughly *M* by *N*. The first array is the positional prefix array, *a*. The positional prefix array has size *M* × (*N* + 1), it has *N* + 1 columns of height *M*. Column *i* (*a*[*i*]) contains the strings in *X* sorted by their reversed prefixes of length *i*. If two strings have the same reversed prefixes of length *i*, their relative order is the same in *a*[*i*] and *a*[0]. *a*[*i*][0] contains the string with the lexicographically smallest reversed prefix of length *i* and *a*[*i*][*N* − 1] contains the string with the lexicographically largest reversed prefix of length *i*. The divergence array, *d*, is size *M* × (*N* + 1). *d*[*i*][*j*] contains the starting position of the longest match between *a*[*i*][*j*] and *a*[*i*][*j* − 1] that ends at position *i* − 1 (inclusive). If no such match exists, *d*[*i*][*j*] = *i*. The last two arrays are the *u* and *v* arrays, each of size *M* × *N*. These arrays keep track of the position *a*[*i*][*j*] would have in *a*[*i* + 1] if it had 0 and 1 at index i respectively. I.E. *u*[*i*][*j*] = the position in *a*[*i* + 1] that *a*[*i*][*j*] would have if it had a 0 at index *i*. *v*[*i*][*j*] = the position in *a*[*i* + 1] that *a*[*i*][*j*] would have if it had a 1 at index *i*.

In this paper, we will use a variation of the PBWT that allows an alphabet of arbitrary size. The key difference is that instead of using the *u* and *v* arrays to keep track of next position, we use a three dimensional array, *w*, of size *M* × *N* × |*Σ*|, where *Σ* is the alphabet. *w*[*i*][*j*][*c*] holds the position *a*[*i*][*j*] would have in *a*[*i* + 1] if it had *c* at index *i*. This variation of the PBWT uses *O*(*MN*|*Σ*|) space instead of the previous *O*(*MN*). However, the set maximal match query time complexity remains the same as before (*O*(*N* + *c*) where *c* is the number of matches found). This variation of the PBWT has been explored by Naseri et al. [10].

We use Durbin’s definitions of local and set maximal matches [6]. This description is from [13]. A match between two strings *s* and *t*, (*i, j, s*) = (*i,j,t*) is **locally maximal** if it can’t be extended in any direction and still match. (*s*[*i* – 1] ≠ *t*[*i* – 1] or *i* = 0) and (*s*[*j* + 1] ≠ *t*[*j* + 1] or *j* = *N* – 1). For a string *s* ∉ *X*, a match between *s* and *x* ∈ *X* ((*i, j, s*) = (*i, j, x*)) is **set maximal** from *s* to *X* if there does not exist a match from *s* to a string in *X* that is larger and contains this match. ∀*t* ∈ *X*∀*k* ∈ {0, …, *i*} ∀*l* ∈ {*j*, …, *M*}, (*k* = *i* and *l* = *j*) or *t*[*k, l*] ≠ *s*[*k, l*].

### 2.2 Minimal Positional Substring Cover (MPSC)

Sanaullah et al. formulated the haplotype threading as the Minimal Positional Substring Cover problem [13]. The Minimal Positional Substring Cover (MPSC) problem is, given a query string *z* and set of strings *X*, find a positional substring cover of *z* by *X* with the smallest size of all positional substring covers of *z* by *X*.

Sanaullah et al. provided an algorithm that outputted an MPSC of *z* by *X* in *O*(*N*) time given a PBWT of *X*, where *N* is the length of *z*. They also provided algorithms for outputting leftmost, rightmost, and set maximal match only MPSCs in *O*(*N*) time given a PBWT of *X*. Lastly, they provided an algorithm that outputted an *h*-MPSC of *z* by *X* in 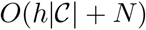 time, where 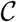 is the outputted *h*-MPSC.

A leftmost MPSC of *z* by *X* is an MPSC where each *i*-th positional substring starts as early as the *i*-th positional substring of every MPSC of *z* by *X*. Similarly, a rightmost MPSC of *z* by *X* is an MPSC where every *i*-th positional substring ends as late as every *i*-th positional substring in an MPSC of *z* by *X*. A set maximal match only MPSC of *z* by *X* is an MPSC of *z* by *X* such that every positional substring it contains corresponds to a set maximal match from *z* to *X*. Lastly, an *h*-MPSC of *z* by *X* is the smallest positional substring cover of *z* by *X* such that all of the positional substrings it contains are contained in *h* strings in *X*. By the definitions on positional substring covers, the size of an MPSC is the number of positional substrings it contains and the length of an MPSC is the sum of the lengths of its positional substrings.

## 3 Methods

### 3.1 MPSC Solution Space

Although a minimal positional substring cover of a query haplotype *z* by a panel *X* is an inference for haplotype threading, there may be many possible minimal positional substring covers of *z* by *X*. The MPSC outputted may not be the most accurate haplotype threading. Therefore, we consider the set of all possible MPSCs of *z* by *X*.

We begin the exploration of the solution space of minimal positional substring covers of *z* by *X* by attempting to bound it. We already have two useful bounds that can be efficiently obtained, namely the leftmost and rightmost MPSCs. The starting point of every *i*-th positional substring in an MPSC of *z* by *X* must be at least 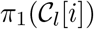, the starting point of the *i*-th positional substring of 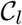, the leftmost MPSC. Likewise, the ending point of every *i*-th positional substring in an MPSC of *z* by *X* is at most 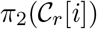, the ending point of the *i*-th positional substring of 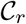, the rightmost MPSC. Sanauallah et al. proved the existence of leftmost and rightmost MPSCs for any *z* and *X* where there exists an MPSC of *z* by *X* [13]. Formally, an MPSC 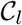 is leftmost if 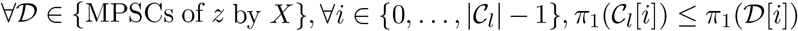. While an MPSC 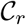 is rightmost if 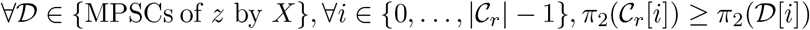.

We have even tighter bounds of the solution space of MPSCs. Sanaullah et al. showed that if the *i*-th positional substring in a leftmost MPSC of *z* by *X* begins at index *j*, every *i* – 1-th positional substring in an MPSC of *z* by *X* contains the index *j* − 1 [13]. Here, we show a similar property for rightmost MPSCs.

#### Claim 1.

Every *i* + 1-th positional substring in any MPSC of *z* by *X* contains index 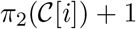. Where 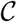 is a rightmost MPSC of *z* by *X*.

*Proof*. Suppose there existed an 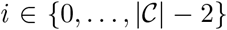 and 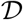, MPSC of *z* by *X*, s.t. 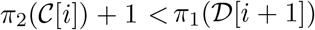. Then, since 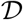 is a cover of *z* by *X*, it contains a positional substring that contains index 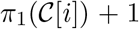, call it 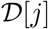. If *j* < *i* + 1, then by definition of *i*-th positional substring, 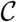 is not rightmost since 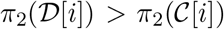. If *j* > *i* + 1, the definition of *i*-th positional substring is contradicted since 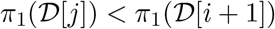 and *j* > *i* + 1. Therefore, no such *i* and 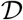 exist.

Suppose there existed an 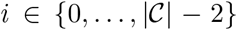 and 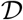, MPSC of *z* by *X*, s.t. 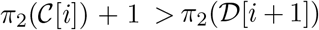. Then, the set 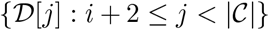 covers the indices 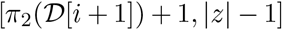 with 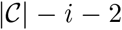 positional substrings and the set 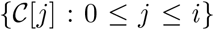 covers the indices 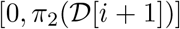 with *i* + 1 positional substrings. Their union is a positional substring cover of *z* by *X* with 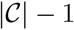 positional substrings. This contradicts the fact that 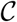 is an MPSC of *z* by *X*. Therefore, no such *i* and 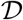 exist. □

For every *i*-th positional substring in a minimal positional substring cover of *z* by *X*, we now have two positions that it is guaranteed to contain: 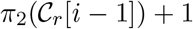 + 1 and 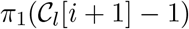. Where 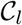 is a leftmost MPSC of *z* by *X* and 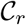 is a rightmost MPSC of *z* by *X*. Furthermore, there are MPSCs of *z* by *X* where the positions just after and before these positions are not contained in the *i*-th positional substrings (rightmost and leftmost MPSCs of *z* by *X* with no overlap, respectively). Therefore, we have the exact range of sites common to all *i*-th positional substrings in MPSCs of *z* by *X*. Call it the *i*-th **required region**. The *i*-th required region is contained in every *i*-th positional substring in an MPSC of *z* by *X*. The *i*-th required region is exactly the first site after the *i* – 1-th positional substring in a rightmost MPSC to the last site before the *i* + 1-th positional substring in a leftmost MPSC. See Fig. 1 for a depiction of required regions and how they are obtained. The *i*-th required region is defined and its properties proven in Lemma 1.

#### Lemma 1 Required Regions

*There exists a contiguous nonempty range of sites for every* 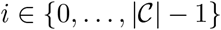, *such that* 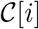 *contains this range of sites for all MPSCs* 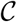 *of z by X. Call the largest such range the i-th required region. For* 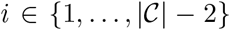, *this range is* 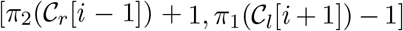, *where* 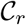 and 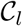 *are rightmost and leftmost MPSCs of z by X respectively. The required region for* 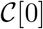 *is* 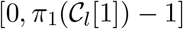, *and for* 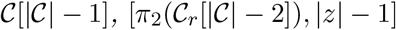, |*z*| − 1].

*Proof*. By Claim 1, every *i*-th substring must contain index 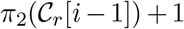 for 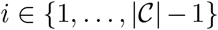. The 0-th substring must contain index 0 by definition of MPSC and *i*-th substring (every site is covered and 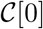 has the smallest starting point of all positional substrings in 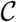). Therefore, the required region exists and is nonempty for all *i*.

In their Claim 5, Sanaullah et al. showed that every *i*-th substring must contain index 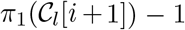 for 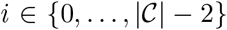 [13]. Furthermore, the 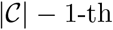 positional substring must contain index |*z*| − 1 by definition of MPSC and *i*-th substring (every site is covered and 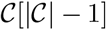 has the largest ending point of all positional substrings in 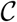).

Positional substrings cover a contiguous range of sites. Therefore the *i*-th required regions for 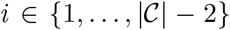 contains indices 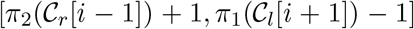. 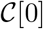 and 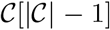 must contain indices 0 and |*z*| − 1 respectively by definition of MPSC and *i*-th positional substring.

Lastly, these ranges are the complete required regions. For 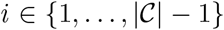, there exists an MPSC in which the *i*-th positional substring doesn’t contain index 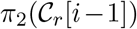, namely a rightmost MPSC with its *i*-th positional substring trimmed to only include sites not covered by its *i* − 1-th substring. For 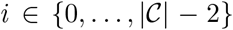, there exists an MPSC in which the *i*-th positional substring doesn’t contain 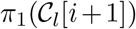, namely a leftmost MPSC with its *i*-th positional substring trimmed to only include sites not covered by its *i* + 1-th substring. Finally, −1 is not part of the 0-th required region and |*z*| is not part of the 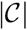 −1-th required region because it is impossible for either of these indices to be contained in an MPSC of *z* by *X*. □

**Fig. 1.**
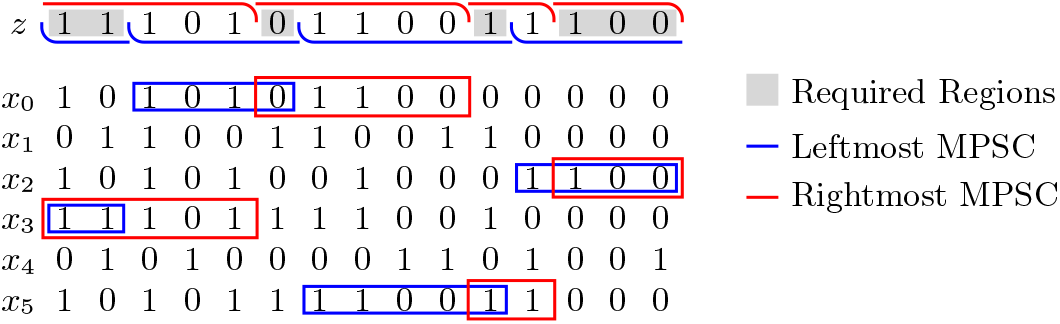
Required regions for MPSC of *z* by *X*. The *i*-th required region is bounded by the *i* + 1-th positional substring in the Leftmost MPSC and the *i* – 1-th positional substring in the Rightmost MPSC.

#### MPSC Graph

We attempt to represent the solution space of MPSCs as a graph. Positional substrings are vertices in the graph and edges occur between positional substrings that are adjacent or overlapping. Consider the naive graph *G* = (*V, E*) where the set of nodes *V* is the set of positional substrings that are contained in both *z* and *X* with *z* as the source string. I.E. *v* = (*i, j, z*) ∈ *V* ⇔ (*i, j, z*) is present in *z* and *x* ∈ *X*. There is an edge between two positional substrings *u* and *v* if *u* starts before *v* and *u* and *v* are adjacent or overlapping (*π*_1_(*u*) ≤ *π*_1_(*v*) and *π*_2_(*u*) ≥ *π*_1_(*v*) − 1). In this graph, all shortest paths from (−1, 0, *z*) to (*N* −1, *N, z*) correspond to minimal positional substring covers of *z* by *X*. Furthermore, all minimal positional substring covers are represented by a shortest path from (−1, 0, *z*) to (*N* −1, *N, z*). However, there are possibly *O*(*N*^2^) nodes and *O*(*N*^4^) edges in this graph, so a shortest path finding algorithm would run in *O*(*N*^4^) time. Therefore, we attempt to simplify the graph. We begin with the following observations.

#### Claim 2.

Every non-empty positional substring *m* = (*i, j, z*) contained in *z* and a string in *X* is contained in a set maximal match from *z* to *X*.

*Proof*. If *m* is a set maximal match, it is contained in a set maximal match (itself). If *m* is not a set maximal match, then there exists a larger match, *n*, between *z* and *s* ∈ *X* that contains *m*. If *n* is set maximal, we are done, otherwise, there exists a match larger than n that contains it (and therefore *m*). We can repeat this logic until a set maximal match containing *m* is found. This process is guaranteed to stop because there are a finite amount of matches from *z* to *X* (since *z* and *X* are finite) and each match is considered at most once. □

#### Claim 3.

For any two set maximal matches from *z* to *X, m* = (*i, j, s*) and *n* = (*k, l, t*), *i* = *k* ⇔ *j* = *l*.

*Proof*. For two set maximal matches with the same starting position, if they have different ending positions, the smaller one is contained in the larger one and is not a set maximal match. Similarly, for two set maximal matches with the same ending position, if the have different starting positions, the smaller one is contained in the larger one and is not set maximal. Therefore, two set maximal matches have the same starting position if and only if they have the same ending position. □

Therefore, we consider a graph where the set of nodes is the set of set maximal match positions from *z* to *X*. Every possible non-empty match is contained in at least one of these nodes. There is an edge between nodes *u* and *v* with the same conditions as before: *π*_1_(*u*) ≤ *π*_2_(*v*) and *π*_2_(*u*) ≥ *π*_1_(*v*) − 1. Call the node with index 0 the source node, *s*. Call the node with index *N* − 1 the sink node, *t*. *s* and *t* are unique by Claim 3. There is a one-to-one correspondence between shortest paths from *s* to *t* in this graph and minimal positional substring covers of *z* by *X* composed of set maximal matches. Furthermore, there is a one-to-one correspondence between shortest paths in the graph from *s* to *t* and MPSCs of *z* by *X*. The *i*-th positional substring in the MPSC is contained in the *i*-th node in the path it maps to. By Claim 3, the number of nodes in this graph is *O*(*S*) ⊆ *O*(*N*). However, the number of edges in this graph may be *O*(*S*^2^), therefore a shortest path finding algorithm on this graph would still take *O*(*S*^2^) ⊆ *O*(*N*^2^) time. We simplify the graph of the solution space again, this time considering the *i*-th required regions.

We will construct a graph where the set of nodes is the set of set maximal match positions from *z* to *X*. However, we will also consider the information gained from the *i*-th required regions. For two nodes *u, v*, there is an edge from *u* to *v* in the graph if the previous requirements are fulfilled, *u* contains the complete *i*-th required region, and *v* contains the complete *i* + 1-th required region. The following property is useful in this construction.

#### Claim 4.

For every positional substring that is contained in *z* and a string in *X*, if it contains a full required region, it doesn’t contains sites from any other required region.

*Proof*. If there existed a positional substring *m* that fully contains *i*-th required region and a site from another required region, then *m* must contain sites from either the *i* − 1-th or the *i* + 1-th required region. If it contains sites from the *i* − 1-th required region, we can show that these sites do not belong in the *i* − 1-th required region by replacing 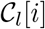 with *m* and removing any overlap of 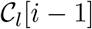 and 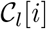 from 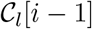 (for leftmost MPSC of *z* by *X*, 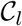).

Similarly, if *m* contains sites from the *i* + 1-th required region, we can show that they are not required by replacing 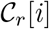 with *m* and removing any overlap between 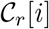 and 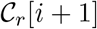 from 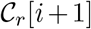 (for rightmost MPSC of *z* by *X*, 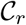). These are contradictions, therefore no such positional substring *m* exists. □

If *u* contains the *i*-th required region and *v* contains the *i* + 1-th required region, then *u* starts before *v*. Therefore, there is an edge from *u* to *v* in this graph iff u contains the *i*-th required region, *v* contains the *i* + 1-th required region, and *u* and *v* are adjacent or overlapping. See Fig. 2 for a depiction of this graph for *z* and *X*. All paths from *s* to *t* in this graph correspond to a MPSC of *z* by *X* composed of only set maximal matches. Furthermore, all MPSCs of *z* by *X* correspond to a path from *s* to *t* in the graph where the *i*-th positional substring is contained in the *i*-th set maximal match position in the path.

**Fig. 2.**
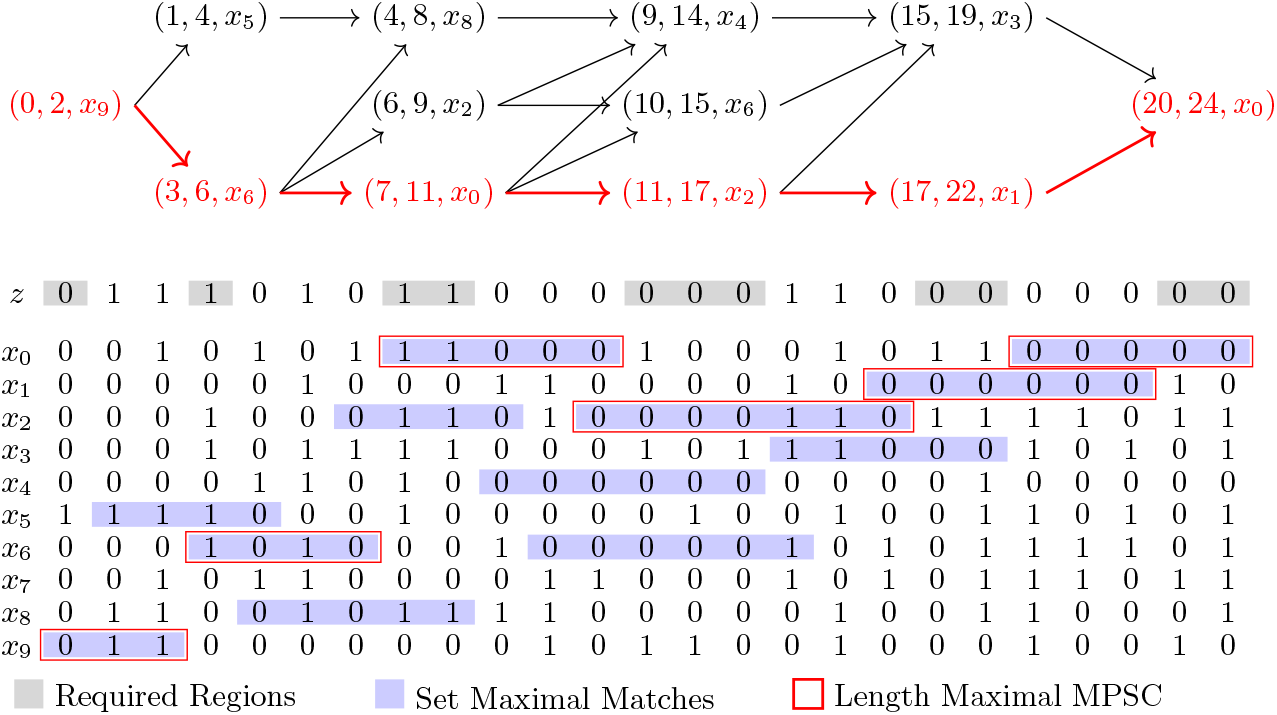
Constructed MPSC graph of *z* by *X*. In gray are the required regions for *z* and *X*. Highlighted in blue are the set maximal matches from *z* to *X*. Highlighted in red is the Length Maximal MPSC of *z* by *X*.

This graph can be constructed very efficiently. As before, there are *O*(*S*) ⊆ *O*(*N*) nodes in the graph. These nodes can be found in *O*(*N*) time. All *i*-th required regions can be obtained in *O*(*N*) time. This is done by first calculating a leftmost MPSC and a rightmost MPSC of *z* by *X* in *O*(*N*) time [13]. Then, each *i*-th required region can be obtained in constant time by Lemma 1. The set maximal match positions can be obtained in *O*(*N*) time [14]. The last step is the creation of the edges between nodes. Although there may be *O*(*S*^2^) ⊆ *O*(*N*^2^) edges in this graph, we avoid the explicit construction of all of them by exploiting the following property. Call *R_i_* the set of nodes that contained the *i*-th required region. Call *R_i_*[*j*] the node in *R_i_* with the *j*-th smallest starting position, *j* ∈ { 0, …, |*R_i_*| − 1}. Then,

#### Lemma 2.

*The set of nodes R_i_*[*j*] *has an edge to is a subset of the set of nodes R_i_*[*j* + 1] *has an edge to*.

*Proof*. All of the nodes *R_i_*[*j*] has an edge to are in *R*_*i*+1_. The set maximal match of every node in *R*_*i*+1_ starts after the *i*-th required region by Claim 4. *R_i_*[*j*] ends before *R_i_*[*j* + 1], and both start at or before the start of the *i*-th required region. Therefore, every set maximal match in *R*_*i*+1_ that overlaps or is directly after *R_i_*[*j*] overlaps *R_i_*[*j* + 1]. Therefore, the set of nodes *R_i_*[*j*] has an edge to is a subset of the set of nodes *R_i_*[*j* + 1] has an edge to. □

Therefore, during the construction of our graph, we don’t directly construct all *O*(*N*^2^) edges. For every node *R_i_*[*j*+1], we only construct edges to nodes that *R_i_*[*j*] does not have an edge to. Although, there may be *O*(*N*^2^) edges in the complete graph, we fully encode them with *O*(*S*) ⊆ *O*(*N*) edges. With these edges in place and the property in Lemma 2 in mind, we have constructed a graph that fully describes the solution space of MPSCs of *z* by *X* in *O*(*N*) time. The combination of this graph and the PBWT supports many efficient queries through the exploitation of Lemma 2. This includes the counting of the number of set maximal match only MPSCs of *z* by *X*, the counting of the number of MPSCs a positional substring is an element of, and the counting of Length Maximal MPSCs, all in *O*(*N*) time. Furthermore, each of these sets of MPSCs can be enumerated in *O*(*N* + *S_c_*) time, where *S_c_* is the number of MPSCs outputted. These problems are very useful for the efficient and accurate imputation and phasing of haplotypes.

### 3.2 Length Maximal MPSC

The Length Maximal Minimal Positional Substring Cover problem is, given a set *X* of *M* strings of and a string *z*, find a MPSC of *z* by *X* that has the largest length out of all MPSCs of *z* by *X*. Note that by Claim 2 of [13], the length of any MPSC of *z* by *X* is less than or equal to 2*N*.

#### Lemma 3.

*Every positional substring in any Length Maximal MPSC of z by X is a set maximal match from z to X*.

*Proof*. Suppose there existed a length maximal MPSC of *z* by *X*, 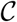, that contained a positional substring that is not a set maximal match. Then, replacing it with a positional substring that is a set maximal match that contains it yields another MPSC 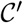 with a larger length. Such a positional substring always exists by Claim 2. 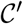 is an MPSC because it covers all the sites 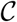 covered and has the same size. Its length is larger because the only difference between 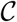 and 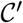 is a positional substring in 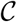 was replaced by a larger one 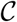. This contradicts the fact 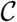 is a length maximal MPSC of *z* by *X*. Therefore, every length maximal MPSC of *z* by *X* is composed of only set maximal matches.

Considering Lemma 3, we begin with the MPSC graph in Fig. 2 and modify it in the following fashion. For every edge (*u, v*), we assign it a weight of the length of *u*. In this directed acyclic graph, every directed path from *s* to *t* still corresponds to a MPSC of *z* by *X* composed of only set maximal matches. As before, every MPSC of *z* by *X* composed of only set maximal matches corresponds to a directed path in the graph from *s* to *t*, there is a one-to-one correspondence. Therefore, the length maximal MPSC of *z* by *X* corresponds to a path in the graph by Lemma 3. In fact, there is a one-to-one correspondence between longest directed paths in the graph from *s* to *t* and Length Maximal MPSCs of *z* by *X* (where the length of a path is the sum of the weights of its edges).

Therefore, we can obtain a Length Maximal MPSC by constructing this graph and finding a longest path in it from *s* to *t*. This is easy given a PBWT of *X*. We begin by finding leftmost and rightmost MPSCs of *z* by *X*. This can be done in *O*(*N*) given a PBWT of *X* (where *N* is |*z*|). This was shown by Sanaullah et al. [13]. We can then obtain all required regions in *O*(|*C*|) time, where |*C*| is the size of an MPSC of *z* by *X*. The calculation of each required region takes constant time. They can be calculated simply using the leftmost and rightmost MPSCs and Lemma 1. Next, we output all set maximal match positions and one string in *X* that each contains each set maximal match in *O*(*N*) time. This can be done with a simple modification of the set maximal match query algorithm by Sanaullah et al. [14]. We maintain the sorted order of the set maximal match positions provided by the query algorithm. Lastly, we keep track of which set maximal matches contain which complete required regions. This can be done in a simple sweeping fashion in *O*(*S*) time due to Claim 4. So far, the algorithm has taken *O*(*N*) time.

We now have all the required information to build the graph. We create the nodes of the graph in *O*(*S*) time where *S* is the number of set maximal match positions, *S* ≤ *N*. We create the edges leaving *s* and entering *t* in *O*(*S*) time. Lastly, for every set maximal match position, if it fully contains a required region, we create an edge from it to nodes whose positional substrings fully contain the next required region and are adjacent or overlapping. This last step of construction of the graph takes *O*(*S*^2^) ⊆ *O*(*N*^2^) time. We can find the longest path in this directed acylic graph in *O*(*V* + *E*) = *O*(*S* + *E*) ⊆ *O*(*S*^2^) ⊆ *O*(*N*^2^) time. This can be done using a shortest path finding algorithm on the graph with the weights negated. The overall time complexity for this method of outputting a length maximal MPSC is *O*(*N* + *S*^2^). However, we have yet to exploit the property of the graph shown in Lemma 2.

We will now show that the longest path in the graph can be obtained in time linear to the number of nodes in the graph, *O*(*S*) through the exploitation of Lemma 2. Therefore, a Length Maximal MPSC of *z* by *X* can be outputted in *O*(*N*) time. Call the set of set maximal match positions (or the corresponding nodes) containing the *i*-th required region *R_i_*. The key idea is the following. Given the longest paths from all nodes in *R*_*i*+1_ to *t*, the longest paths from all nodes in *R_i_* to *t* can be obtained in *O*(|*R*_*i*+1_| + |*R_i_*|) time. This is despite the fact that there may be |*R*_*i*+1_| × |*R_i_*| edges between these nodes. We denote the set maximal match of *R_i_* with the *j*-th smallest starting position as *R_i_*[*j*]. I.E. *R_i_*[0] is the set maximal match in *R_i_* with the smallest starting position, *R_i_*[1] has the next smallest starting position, etc. The calculation of the longest paths from *R_i_* to *t* in linear time depends on Lemma 2. The set of nodes *R_i_*[*j*] has an edge to is a subset of the set of nodes *R_i_*[*j* + 1] has an edge to. Given this property, we can calculate the longest paths in a straightforward fashion that evaluates the longest path from each node in *R*_*i*+1_ to *t* at most once. This is done by calculating longest paths of nodes in order of starting position of the corresponding set maximal match, least to greatest. See Fig. 3 for a depiction of this process. Pseudocode of this process is provided in Algorithm 8 in Appendix A.

**Fig. 3.**
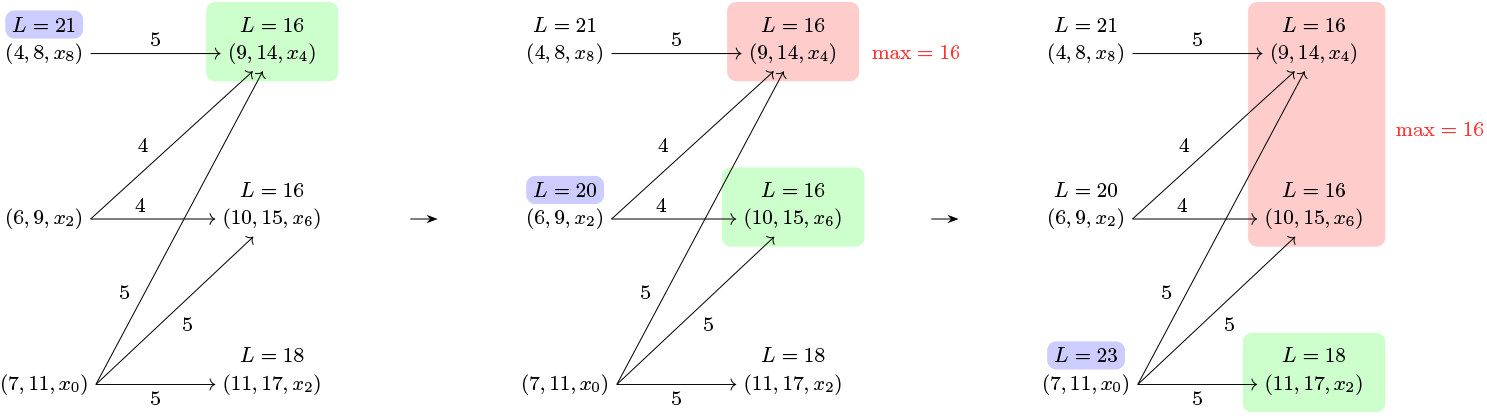
Obtaining the longest paths of the set maximal matches in *R_i_* from the longest paths of the set maximal matches in *R*_*i*+1_ in *O*(|*R_i_*| + |*R*_*i*+1_|) time. Subgraph of Fig. 2, *i* = 2.

After the lengths of the longest paths from each node to *t* is calculated, the longest path from *s* to *t* can be calculated in a simple linear backtracking step. Start with *s*. For each *i*-th required region for 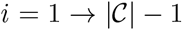, take the first node with length of longest path to *t* equal to the length of the longest path to *t* of the *i* − 1-th node in the path so far minus the length of the substring of the *i* − 1-th node in the path so far. See Algorithm 6 in Appendix A for the pseudocode of this process. With this process for outputting the longest path in the graph from *s* to *t* in *O*(*S*) time, we can find the longest path in the graph without explicitly constructing the graph. Therefore, givena PBWT of *X*, we output a length maximal minimal positional substring cover of *z* by *X* in *O*(*N*) time. The pseudocode of this algorithm can be seen in Algorithm 1 in Appendix A.

### 3.3 *h*-MPSC

An *h*-MPSC of a query string *z* by a set of strings *X* is a smallest set of positional substrings that forms a positional substring cover of *z* by *X* and are each contained in at least *h* strings in *X*. An MPSC of *z* by *X* is an *h*-MPSC of *z* by *X* where *h* = 1. Sanaullah et al. provided an algorithm for outputting an *h*-MPSC, 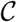, of *z* by *X* in 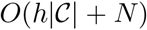 time given a PBWT of *X* [13]. We suspect their algorithm outputs a “leftmost” *h*-MPSC. Here we describe an algorithm that improves on the previous h-MPSC algorithm running time. It outputs an *h*-MPSC of *z* by *X* in *O*(*N*) time. We believe the *h*-MPSC it outputs is “rightmost”.

The original *h*-MPSC algorithm traversed the PBWT from index *N* to 0. Here, we traverse in the opposite direction, from 0 to *N*. We begin at index 0 and keep track of the block of strings that match with *z* on the range [*i, j*). The block is denoted by [*f, g*) where *f* is the first string in the block in the prefix sorting and *g* is the first string not in the block after *f*. If *f* = *g*, the block is empty. At site 0, *f* and *g* are initialized to *f* = 0, *g* = *M*. *i* and *j* are initialized to 0. The *f* and *g* blocks are updated in constant time per site using the *w* array between sites. Once the block at site *j* + 1 has less than *h* strings, the positional substring of the previous block (i, *j* −1, *z*) is added to the *h*-MPSC. This is repeated until all sites are covered by a positional substring contained in *h* strings in *X*. This algorithm runs in *O*(*N*) time because the is traversed from index 0 to *N* and each site takes constant time to update the *f* and *g* block. Furthermore, the addition of a positional substring to the *h*-MPSC can happen at most once per site and takes constant time. A proof that the positional substring cover is an *h*-MPSC can be seen through a fairly simple modification of Lemma 2 in [13]. The pseudocode of this algorithm can be seen in Algorithm 9 in Appendix B.

## 4 Results

### 4.1 Haplotype Threading Properties

We explored the properties of haplotype threadings of the MPSC formulation. The dataset used was the UK Biobank (UKB) [3]. The UKB has 974,818 haplotypes and around 700,000 sites (microarray). We used chromosome 21, which has 9,793 sites. For each haplotype in the UKB, we run our method to identify an MPSC of it using all other haplotypes in the UKB as the reference panel. We also evaluate a rudimentary method of handling mismatches in the MPSC formulation of haplotype threading by using P-smoother [15]. P-smoother is a method for smoothing out sporadic mismatches in otherwise well-matched PBWT blocks, to attempt to remove very rare mutations and genotyping errors from the British only UKB panel. P-smoother was run on the default settings and flipped the alleles of roughly 1.4% of the data. The smoothed panel is expected to tolerate mismatches, resulting in longer match segments, and smaller MPSC sizes. The benchmarking source code is available at genome.ucf.edu/MPSC for all benchmarks.

We found that the MPSC segment count of an individual has an overall distribution due to the fact that not every one’s relatives are sampled evenly. However, the mode of MPSC segment count is inversely correlated with the number of templates with closely related ethnic background (Fig. 4). This behavior is even more clear when we run the experiment with varied sample sizes (Fig. 5). We plot the number of segments in each haplotype threading by self reported ethnic background in Fig. 4. The x-axis is the number of segments in the haplotype threading and the y-axis is the frequency of that number within each self reported ethnic background. The plotted ethnic backgrounds are the 4 most commonly reported ethnic backgrounds in the UKB: British, Irish, Indian, and Caribbean (860,584, 25,436, 11,320, and 8,598 haplotypes respectively). A haplotype is classified as one of these ethnic backgrounds if it was the first ethnic background the individual who owns the haplotype reported themselves as. We also plot MPSC size distributions for random subsets of the UKB of varying sizes. No threading was found for (7.3%, 0.32%, 0.003%, and 0%) of the haplotypes for M=(1,000, 10,000, 100,000, and 860,022) respectively. *M* = 10 and 100 were also tried, but no haplotype threadings were found in either panel. These haplotypes are left out of the frequency calculation.

**Fig. 4.**
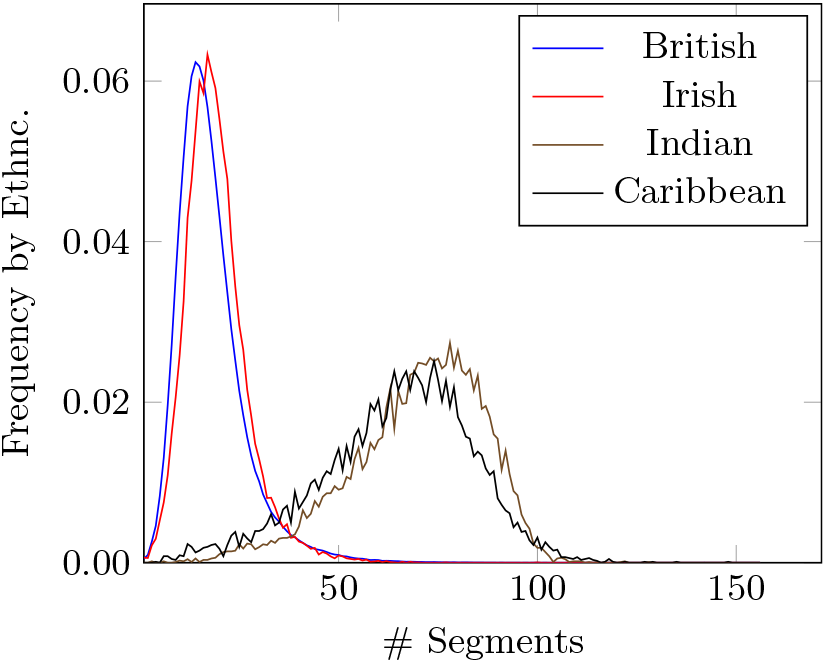
Frequency within self reported ethnic background of # of segments in haplotype threading of haplotypes in UK Biobank. Haplotype threading is generated with reference panel as the rest of the UK Biobank. Frequency is plotted by self reported ethnic background for the four most common in the UK Biobank: British, Irish, Indian, and Caribbean, with 860,584, 25,436, 11,320, and 8,598 haplotypes respectively.

**Fig. 5.**
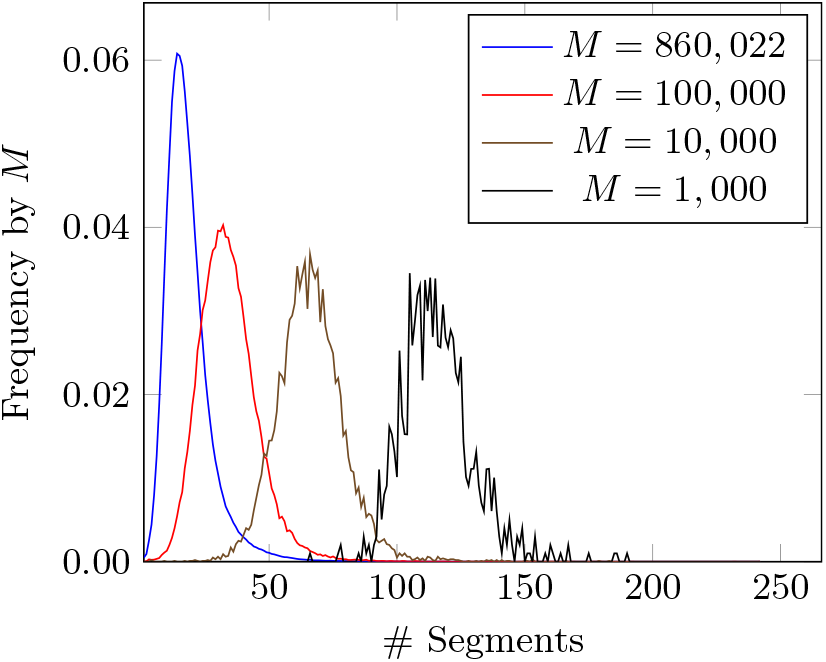
Frequency by panel size of # of segments in British only haplotype threading of haplotypes in UK Biobank. Haplotype threading is generated with reference panel random subset of the British only haplotypes in the UK Biobank. Plotted by panel size for *M* ∈ {1,000, 10,000, 100,000, 860,022}. Note that frequencies exclude haplotypes for which no threading was found.

A more interesting property we consider is the MPSC solution space. This is the number of possible coverings of the query with equally small number of switches, which would be all considered a Viterbi solution for the LS model. We plot the count of Set Maximal Match only MPSCs for each haplotype in the British only UKB dataset (860,022 haplotypes) in Fig. 6 in ascending order. The median solution space size was 120. The 90th and 99th percentile solution space sizes were 72,756 and 3.4 × 10^9^ respectively. As expected, the median solution space size of the P-smoothed panel was 40. The fact that the number of solutions in MPSC solution space is surprisingly small validated that UKB belongs to the regime where MPSC could be a much more efficient alternative for the fully-parameterized Li & Stephens model.

**Fig. 6.**
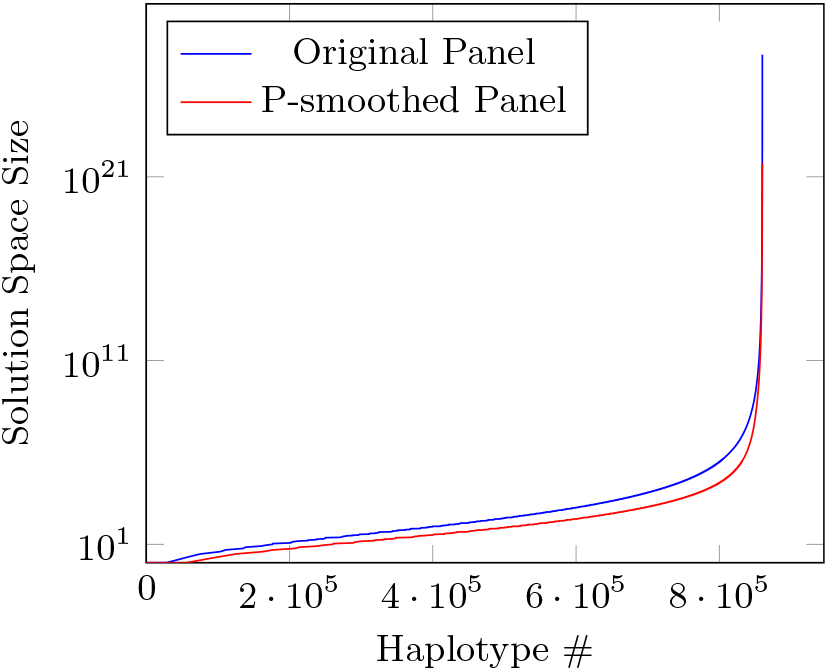
The number of Set Maximal Match only MPSCs per haplotype in British only UK Biobank panel in ascending order.

The last property of the MPSC formulation of haplotype threading we explore is the distributions of lengths of the Length Maximal MPSCs. Note this length *l* is *N* ≤ *l* ≤ 2*N*, and thus the size of the overlap region, *l* – *N* or 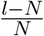, provides one way of quantifying the potential computations (much more than a linear factor!) saved by MPSC compared to enumerating solutions of standard LS. We plot this length distribution of Length Maximal MPSCs for British only haplotypes in Fig. 7. P-smoother results in a smaller Length Maximal MPSC length. The length maximal MPSC provides about at average 1.21X (1.20X after smoothing) coverage of the chromosome, indicating about 20% of the genome belongs to the overlap regions of Length Maximal MPSC where location of recombination breakpoints cannot be unequivocally determined.

**Fig. 7.**
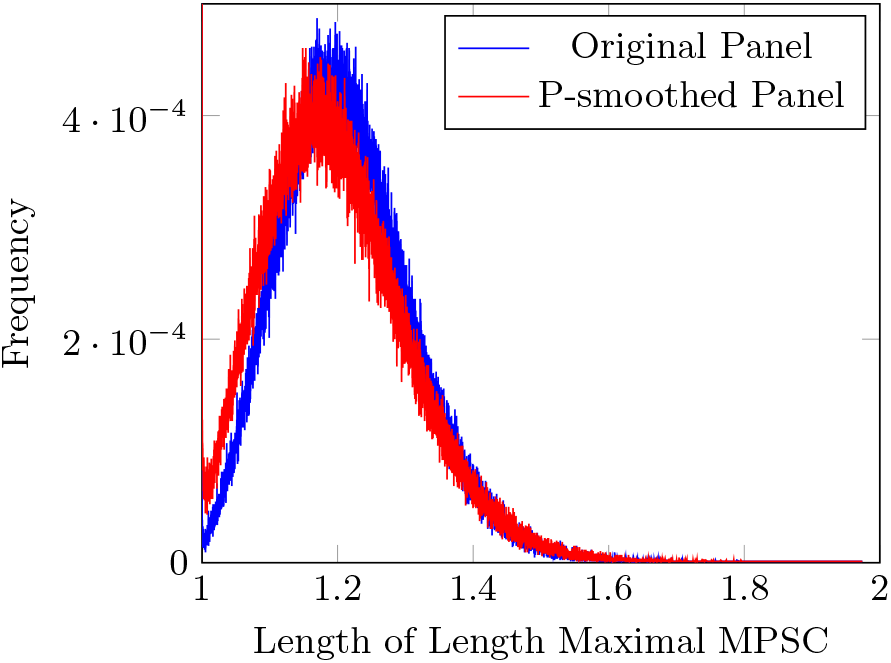
The distributions of lengths of Length Maximal MPSCs for all British only haplotypes in the UKB. Length is represented as a ratio of genome length. The min ratio of an MPSC is 1 and the max ratio is 2. The min and max ratios observed are 1 and 1.974.

### 4.2 Run Time

We measure the run time of obtaining the Length Maximal MPSC for various *M* and *N*. Note that the run time of obtaining a Length Maximal MPSC is strictly larger than the run time of obtaining an MPSC, a Leftmost MPSC, a Rightmost MPSC, and a Set Maximal Match only MPSC. This is because the first three algorithms are subroutines of the Length Maximal MPSC algorithm. The run time of obtaining a Length Maximal MPSC is strictly larger than the run time of obtaining a Set Maximal Match only MPSC because a Length Maximal MPSC is a Set Maximal Match only MPSC. The results of these tests can be seen in Fig. 8.

**Fig. 8.**
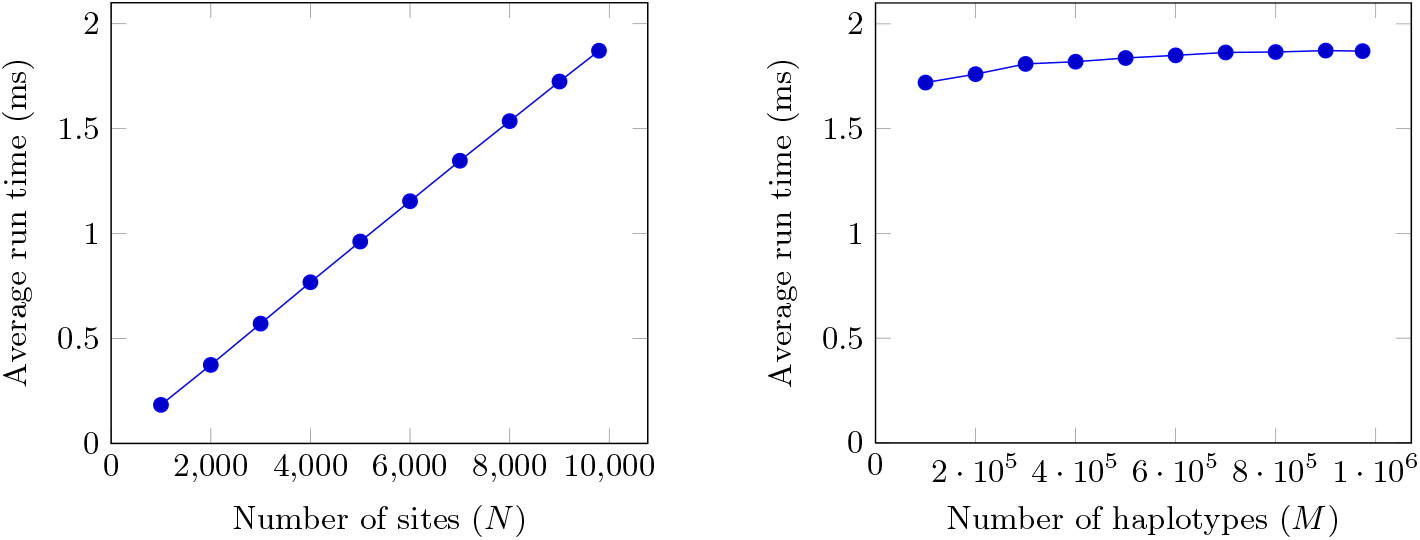
Run time vs varying *M* and *N*. Average run time for the Length Maximal MPSC computation of 1,000 random haplotypes in the UKB British only dataset for varying number of sites, *N* ∈ {1,000, 2,000, 3,000, 4,000, 5,000, 6,000, 7,000, 8,000, 9,000, and 9,793}, is on the left. On the right, average run time for the Length Maximal MPSC computation of 1,000 random haplotypes in the UKB British only dataset for varying number of haplotypes in the reference panel, *M* ∈ {100,000, 200,000, 300,000, 400,000, 500,000, 600,000, 700,00, 800,000, 900,000, and 973,818}, is plotted.

### 4.3 Imputation Benchmark

We implemented an imputation benchmark to demonstrate the usefulness of the MPSC formulation of haplotype threading. The imputation algorithm is naive outside of the haplotype threading: Given a haplotype threading of the haplotype to be imputed at a site, obtain the positional substrings adjacent to it in the haplotype threading. For every haplotype that contains one of these positional substrings, vote once per positional substring it contains towards the allele this haplotype has at the site to be imputed. Impute an allele if the votes are unanimous. We also evaluate the use of a P-smoothed panel. In this method, the haplotype threading is generated using the P-smoothed panel and query haplotype. Votes are counted for haplotypes who have the positional substring in the P-smoothed panel. However, allele votes are counted using the original panel.

This imputation method generalizes to all of the variations of MPSC haplotype threading, therefore we test it on many of the MPSC variations, including Length Maximal MPSC, voting by Set Maximal MPSC Solution Space, and the *h*-MPSC with various choices of *h*. The imputation benchmark was performed in the following fashion: Select 1,000 random haplotypes from the UKB British only dataset (chromosome 21). Remove a random 8,814 sites (90%) from these haplotypes. Impute these haplotypes using the rest of the UKB British only haplotypes (*M* = 859,022) as the reference panel using the above-mentioned MPSC-based imputation methods and Beagle (version 5.4), a state of the art imputation method [2].

As shown in Table 2, all tested MPSC-based imputation methods have an imputation accuracy comparable to Beagle. Interestingly, while MPSC-based methods do not cover all sites, the sites they covered are mostly “easier” sites as Beagle also have higher accuracies on those sites than the average of all sites. Over the covered sites, MSPC-based methods have roughly same accuracies compared to Beagle. Importantly, for a few cases, MPSC-based methods outperform Beagle, especially for Set Maximal MPSC Solution Space (SM MPSC SS), where MPSC methods beat Beagle in both based on the original panel and the P-smoothed panel. This is useful because one can immediately improve the current Beagle results by repainting the Beagle imputation results with MPSC results where MPSC methods cover. Our results also offer a way of studying behaviors of Li & Stephens-based methods. It seems when we increase the *h* in *h*-MPSC, the coverage of MPSC is gravitated towards “easier” regions where both Beagle and MPSC methods have high accuracy. Thus MPSC’s *h* offers a measure of imputation confidence. Finally, the P-smoothed panel increases the power and accuracy of every MPSC threading imputation method.

**Table 2.** Imputation performance in percentages on a random 1,000 British only UKB haplotypes.

| Method | Original Panel |  |  | P-smoothed Panel |  |  |
| --- | --- | --- | --- | --- | --- | --- |
|  | Accuracy |  | Sites Imputed | Accuracy |  | Sites Imputed |
|  | Beagle | MPSC |  | Beagle | MPSC |  |
| Beagle 5.4 | 97.94 |  | 100.00 |  |  |  |
| LM MPSC | 98.45 | 98.43 | 97.15 | 98.43 | **98.46** | 97.23 |
| SM MPSC SS | 99.10 | **99.19** | 88.20 | 99.05 | **99.21** | 88.22 |
| 2-MPSC | 98.70 | 98.32 | 93.14 | 98.67 | 98.36 | 93.32 |
| 3-MPSC | 98.89 | 98.62 | 90.21 | 98.87 | 98.66 | 90.33 |
| 4-MPSC | 99.00 | 98.79 | 87.81 | 98.99 | 98.84 | 87.92 |
| 5-MPSC | 99.12 | 98.96 | 85.71 | 99.10 | 98.99 | 85.86 |
| 8-MPSC | 99.30 | 99.20 | 80.45 | 99.27 | 99.21 | 80.69 |
| 16-MPSC | 99.52 | 99.50 | 71.68 | 99.49 | **99.50** | 72.01 |

Reference panel is the rest of the UKB British only haplotypes. Imputed haplotypes have a random 90% of their sites missing. For each MPSC imputation method, its accuracy and Beagle’s accuracy on the imputed sites are shown. LM MPSC stands for Length Maximal MPSC and SM MPSC SS stands for Set Maximal MPSC Solution Space. MPSC imputation methods that have higher accuracy than Beagle on imputed sites are in bold.

## 5 Conclusion and Discussion

In this paper, we have advanced the MPSC formulation of haplotype threading. We introduced the MPSC graph, a method of representing the solution space of MPSC haplotype threadings of a haplotype that allows many efficient algorithms. These include the enumeration of all MPSCs, Set Maximal Match only MPSCs in optimal time. It also allows the counting of the number of MPSCs a particular positional substring is part of in *O*(*N*) time. Furthermore, we introduced a new MPSC formulation, the Length Maximal MPSC, that maximizes the sum of the lengths of the segments of the haplotype threading over all MPSC haplotype threadings. We provided an algorithm that outputs a Length Maximal MPSC of *z* by a reference panel *X* in *O*(*N*) time given a PBWT of *X*, linear to the number of sites and independent to the number of haplotypes in the panel. We also provided an improved algorithm for the *h*-MPSC problem, where each segment in the threading must be contained in h haplotypes in the reference panel.

Beyond algorithmic contributions, our algorithmic developments established the theoretical basis for linking PBWT and LS-style haplotype threading. When using these algorithms for analyzing properties of MPSC haplotype threadings for the UK Biobank dataset, we demonstrate the usefulness of the MPSC haplotype threadings through an imputation benchmark. We showed that, while the simple MPSC-based imputation does not impute all sites, when it does, the accuracy may be higher than the state-of-the-art imputation method Beagle. Especially, the variations of MPSC algorithms that offer Solution Space, Length Maximal, and high *h* (16) *h*-MPSC can outperform Beagle. In addition, our imputation results verified that P-smoother can be leveraged to soften the mismatch-intolerant MPSC solutions and makes MPSC more robust to real data. We believe further developments based on our sets of algorithms will empower practical applications based on LS such as phasing and imputation.

## Acknowledgments

This work was supported by the National Institutes of Health grants R01 HG010086 and R56 HG011509. This research has been conducted using the UK Biobank Resource under Application Number 24247.

## Appendices

## A Length Maximal MPSC Algorithm

Two versions of the LongestPaths subroutine are provided. The LongestPaths subroutine calculates the length of the longest path from each node to *t* in the graph. There is an in-place and out-of-place version of the subroutine. The out-of-place version is provided because it is easier to understand. Both versions have time complexity *O*(*S*) ⊆ *O*(*N*).

#### Algorithm 1: Output a Length Maximal MPSC of *z* by *X* given a PBWT of *X*

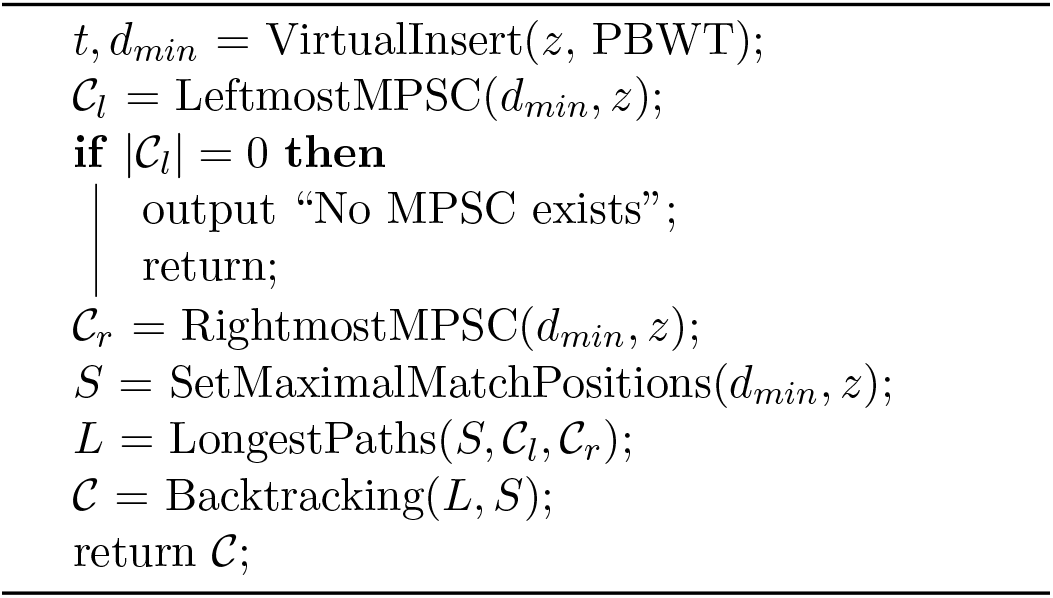

#### Algorithm 2: VirtualInsert: Calculate Prefix and Divergence values of query *z* in a PBWT

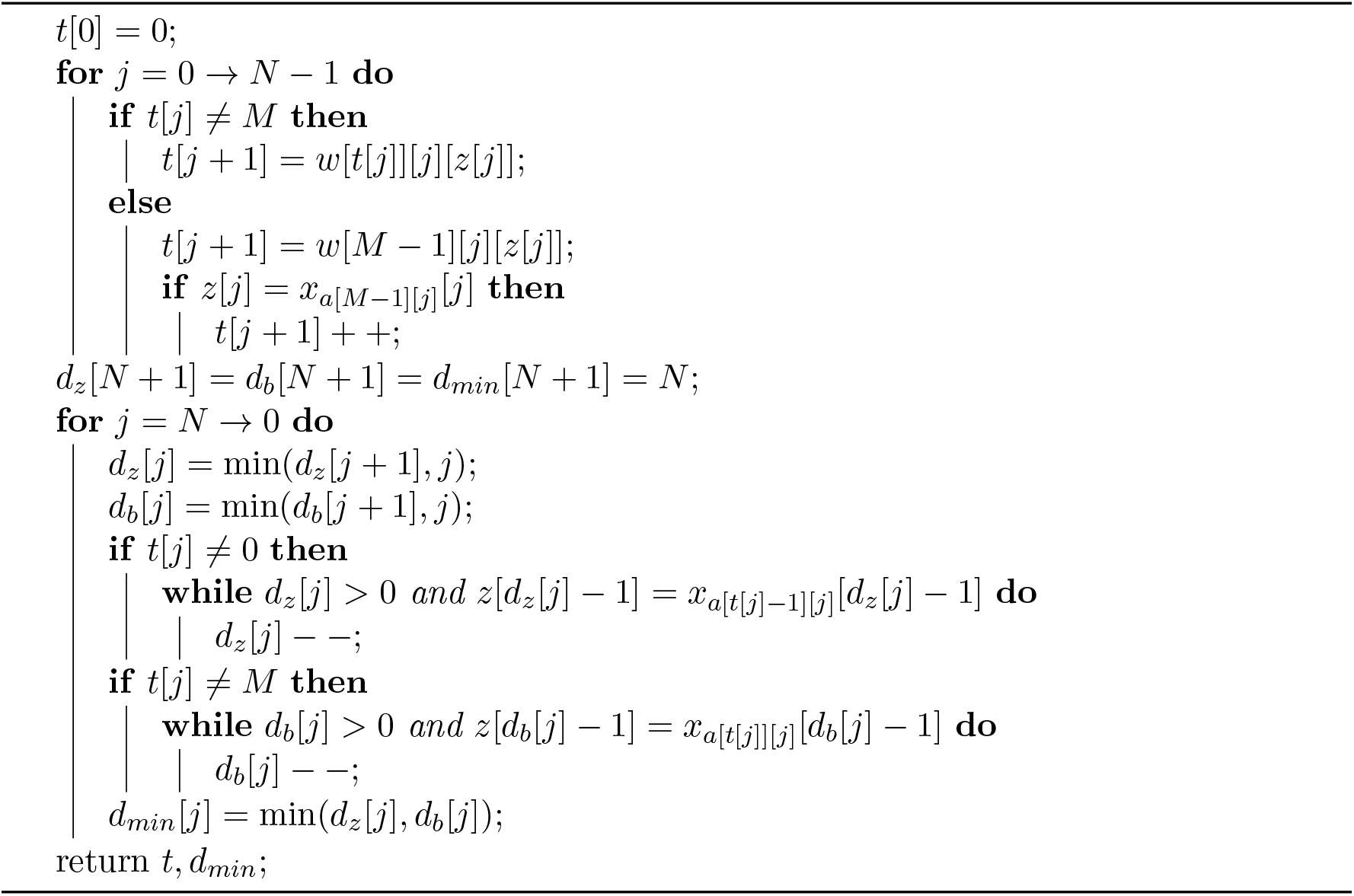

#### Algorithm 3: LeftmostMPSC: Output Leftmost MPSC of *z* by *X*

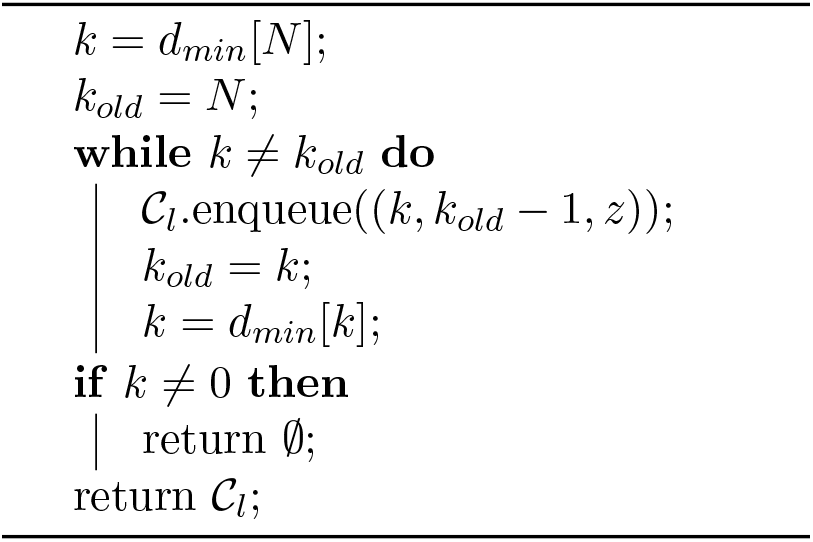

#### Algorithm 4: RightmostMPSC: Output Rightmost MPSC of *z* by *X*

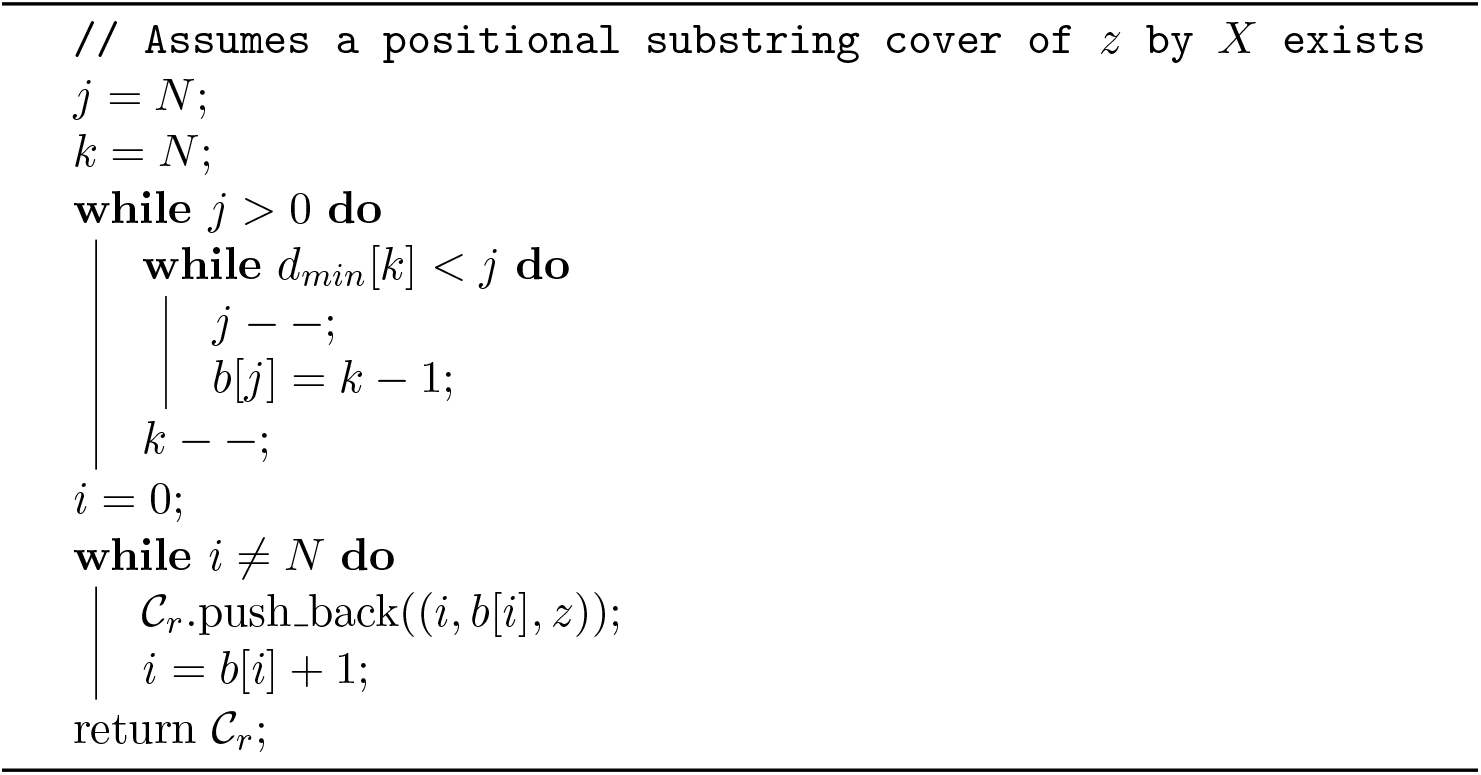

#### Algorithm 5: SetMaximalMatchPositions: Output positions of set maximal matches from *z* to *X*

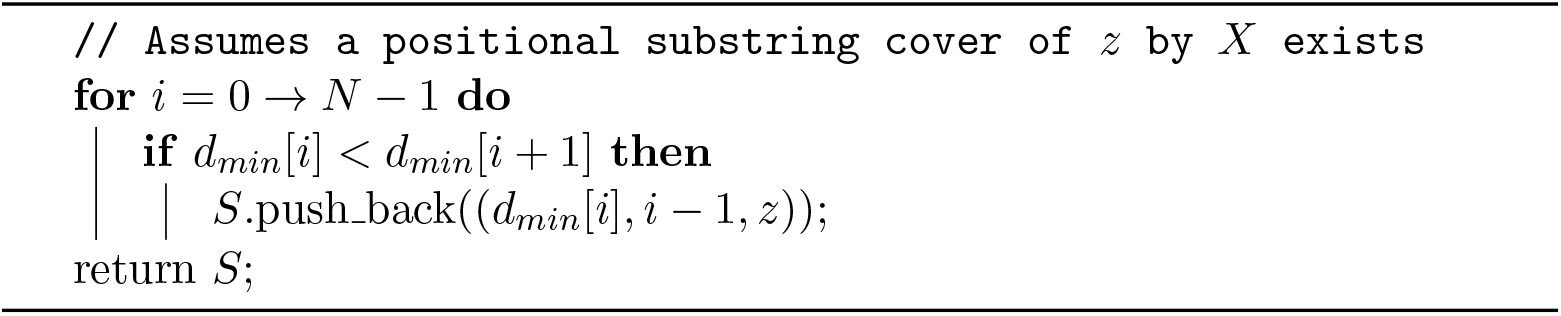

#### Algorithm 6: Backtracking: Output Length Maximal MPSC by backtracking through graph

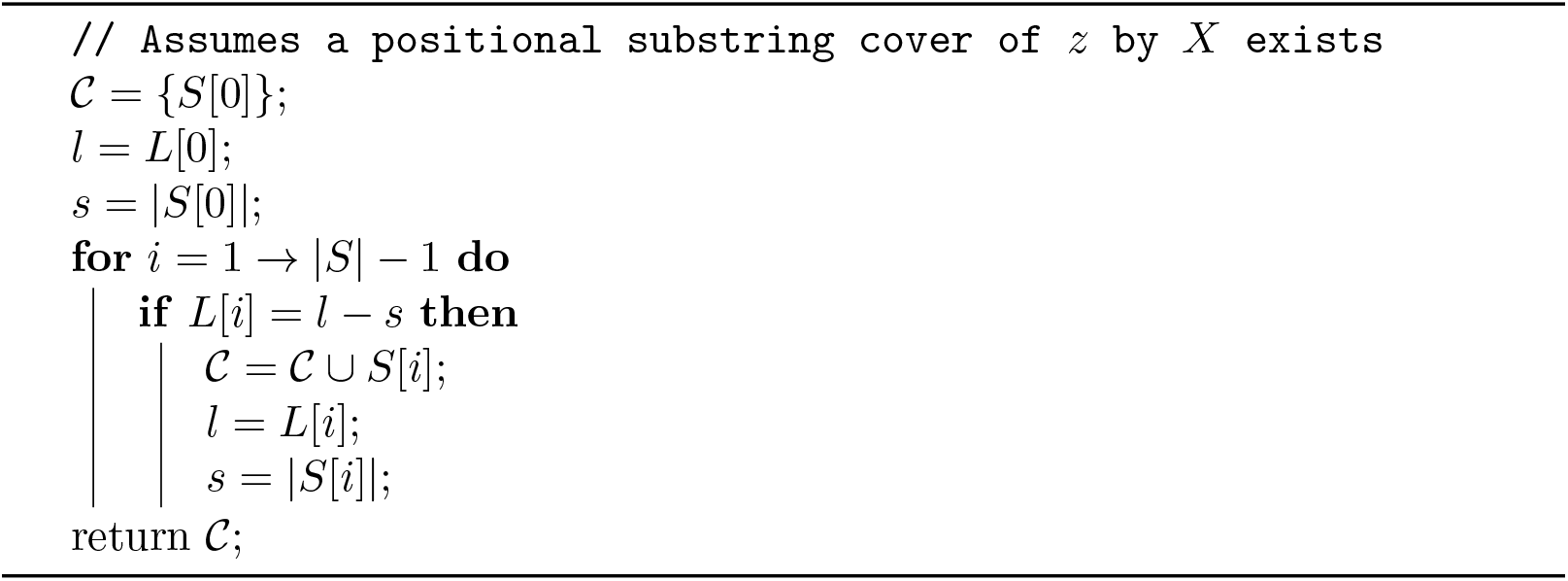

#### Algorithm 7: LongestPaths: Calculate length of Longest Path from every node (out-of-place)

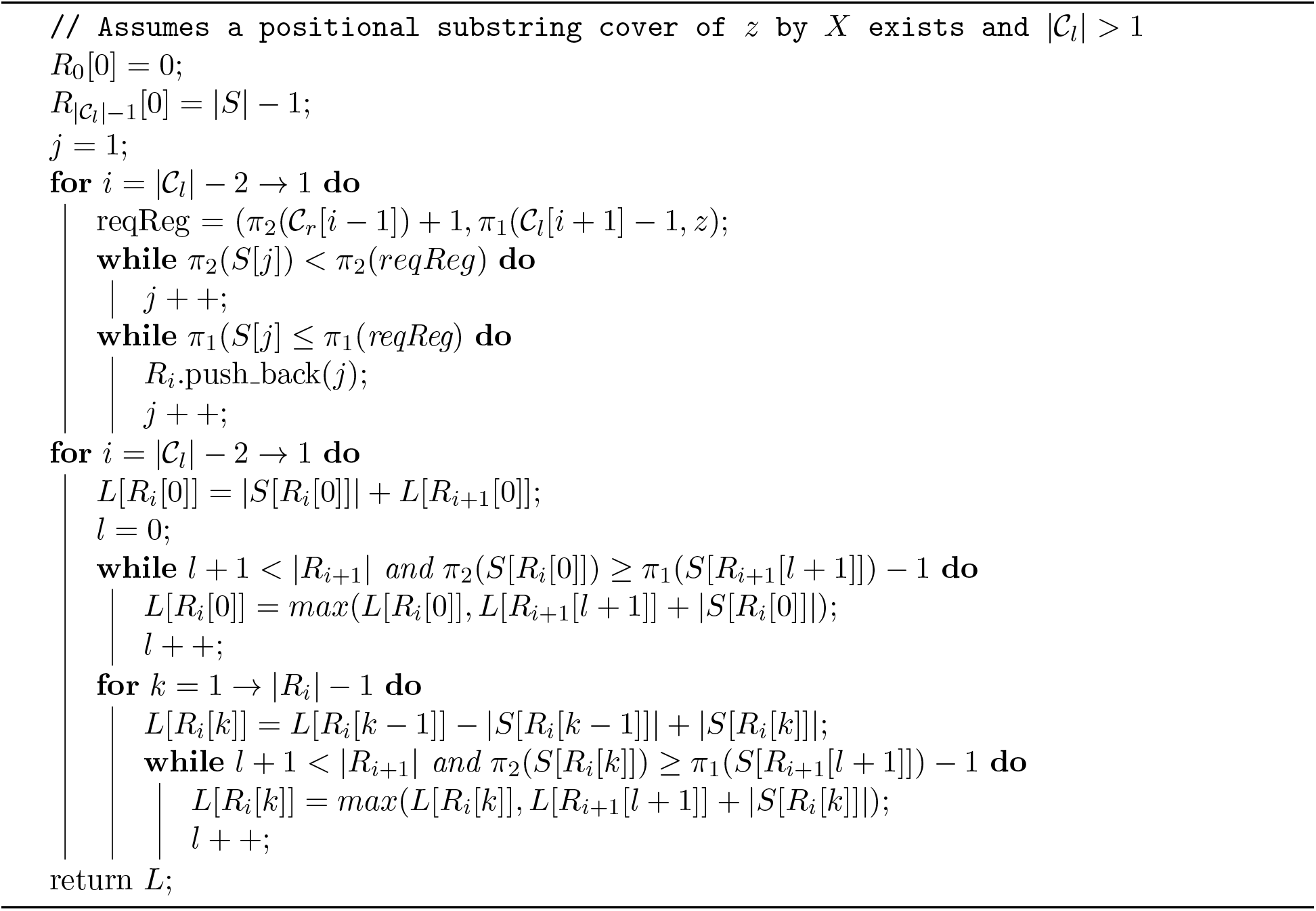

#### Algorithm 8: LongestPaths: Calculate length of Longest Path from every node (in-place)

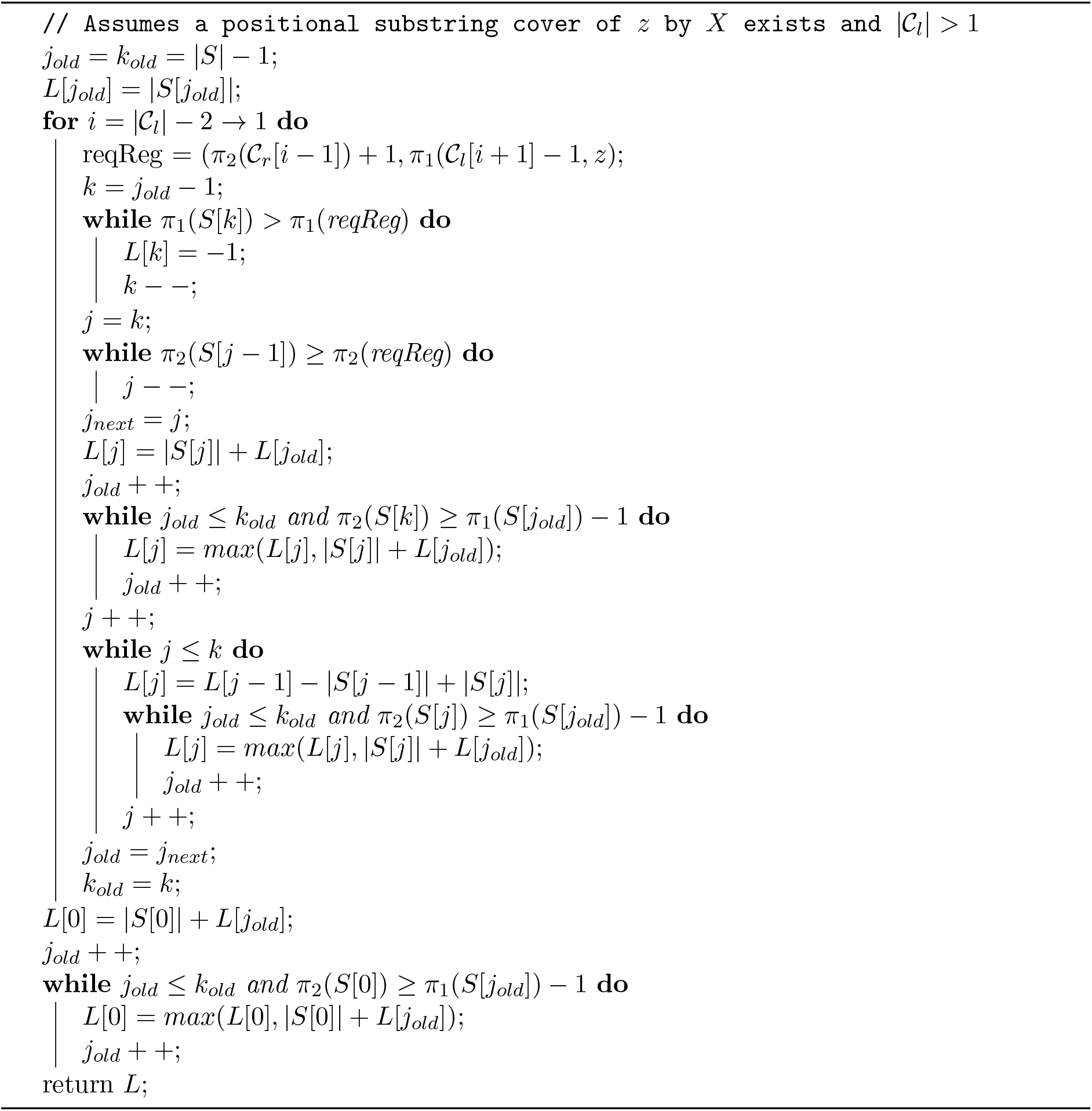

## B *h*-MPSC

### Algorithm 9: *h*-MPSC: Output *h*-MPSC of *z* by *X* in *O*(*N*) time

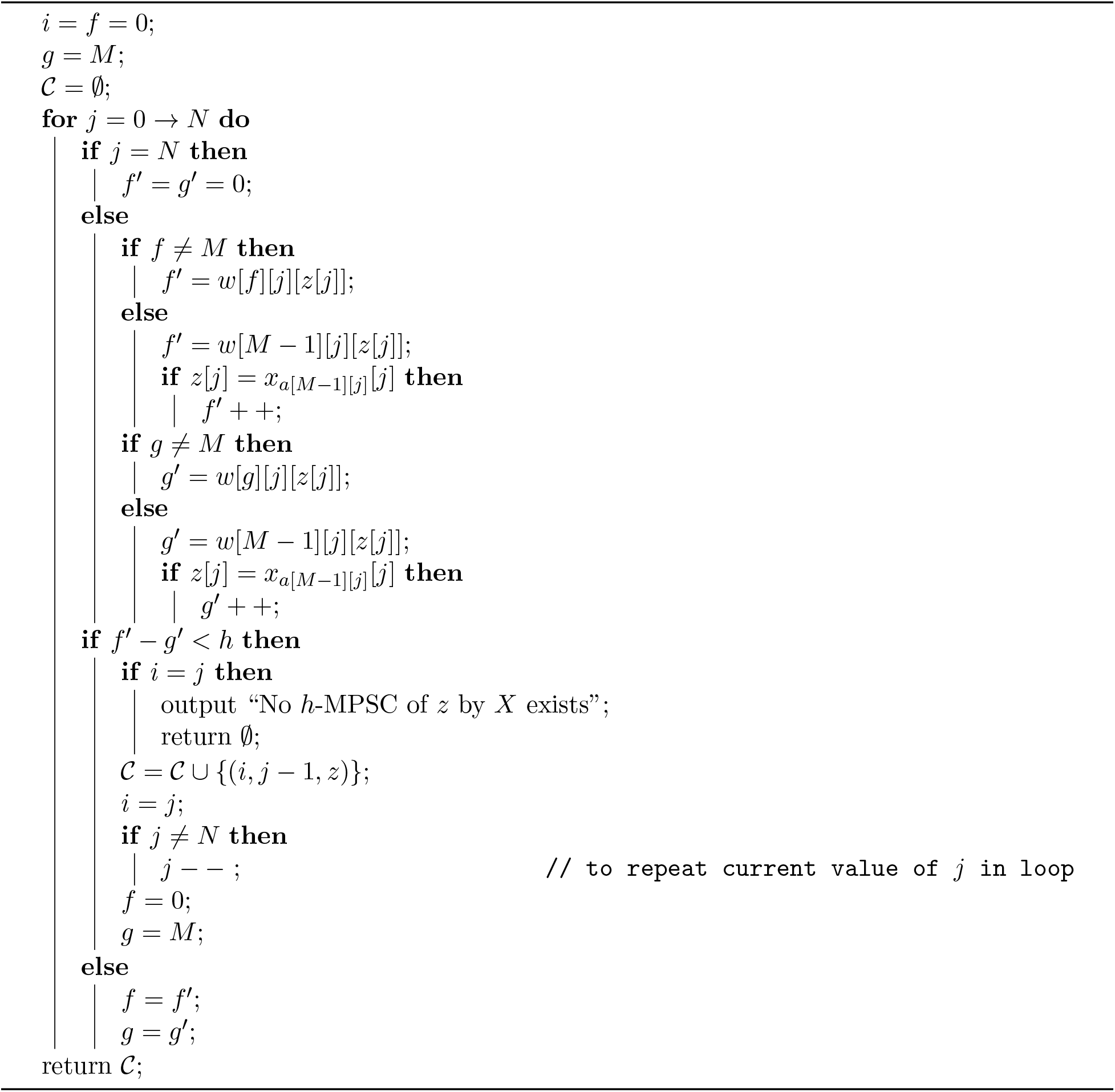

## References

1. Brian L Browning, Xiaowen Tian, Ying Zhou, and Sharon R Browning. Fast two-stage phasing of large-scale sequence data. The American Journal of Human Genetics, 108(10):1880–1890, 2021.

2. Brian L Browning, Ying Zhou, and Sharon R Browning. A one-penny imputed genome from next-generation reference panels. The American Journal of Human Genetics, 103(3):338–348, 2018.

3. Clare Bycroft, Colin Freeman, Desislava Petkova, Gavin Band, Lloyd T Elliott, Kevin Sharp, Allan Motyer, Damjan Vukcevic, Olivier Delaneau, Jared O’Connell, et al. The UK Biobank resource with deep phenotyping and genomic data. Nature, 562(7726):203–209, 2018.

4. Sayantan Das, Lukas Forer, Sebastian Schönherr, Carlo Sidore, Adam E Locke, Alan Kwong, Scott I Vrieze, Emily Y Chew, Shawn Levy, Matt McGue, et al. Next-generation genotype imputation service and methods. Nature Genetics, 48(10):1284–1287, 2016.

5. Olivier Delaneau, Jean-François Zagury, Matthew R Robinson, Jonathan L Marchini, and Emmanouil T Dermitzakis. Accurate, scalable and integrative haplotype estimation. Nature Communications, 10(1):1–10, 2019.

6. Richard Durbin. Efficient haplotype matching and storage using the positional Burrows-Wheeler transform (PBWT). Bioinformatics, 30(9):1266–1272, 2014.

7. Na Li and Matthew Stephens. Modeling linkage disequilibrium and identifying recombination hotspots using single-nucleotide polymorphism data. Genetics, 165(4):2213–2233, 2003.

8. Po-Ru Loh, Petr Danecek, Pier Francesco Palamara, Christian Fuchsberger, Yakir A Reshef, Hilary K Finucane, Sebastian Schoenherr, Lukas Forer, Shane McCarthy, Goncalo R Abecasis, et al. Reference-based phasing using the Haplotype Reference Consortium panel. Nature Genetics, 48(11):1443–1448, 2016.

9. Gerton Lunter. Haplotype matching in large cohorts using the Li and Stephens model. Bioinformatics, 35(5):798–806, 2019.

10. Ardalan Naseri, Degui Zhi, and Shaojie Zhang. Multi-allelic positional Burrows-Wheeler transform. BMC Bioinformatics, 20(11):1–8, 2019.

11. Yohei M Rosen and Benedict J Paten. An average-case sublinear forward algorithm for the haploid Li and Stephens model. Algorithms for Molecular Biology, 14(1):1–12, 2019.

12. Simone Rubinacci, Olivier Delaneau, and Jonathan Marchini. Genotype imputation using the Positional Burrows Wheeler Transform. PLoS Genetics, 16(11):e1009049, 2020.

13. Ahsan Sanaullah, Degui Zhi, and Shaoije Zhang. Haplotype threading using the positional Burrows-Wheeler transform. In 22nd International Workshop on Algorithms in Bioinformatics (WABI 2022). Schloss Dagstuhl-Leibniz-Zentrum für Informatik, 2022.

14. Ahsan Sanaullah, Degui Zhi, and Shaojie Zhang. d-PBWT: dynamic positional Burrows-Wheeler transform. Bioinformatics, 37(16):2390–2397, 2021.

15. William Yue, Ardalan Naseri, Victor Wang, Pramesh Shakya, Shaojie Zhang, and Degui Zhi. P-smoother: efficient PBWT smoothing of large haplotype panels. Bioinformatics Advances, 2(1):vbac045, 2022.

